# Adopting STING agonist cyclic dinucleotides as a potential adjuvant for SARS-CoV-2 vaccine

**DOI:** 10.1101/2020.07.24.217570

**Authors:** Jun-Jun Wu, Yong-Xiang Chen, Yan-Mei Li

## Abstract

A novel STING agonist CDG^SF^ unilaterally modified with phosphorothioate and fluorine was synthesized. CDG^SF^ displayed better STING activity over dithio CDG. Immunization of SARS-CoV-2 Spike protein with CDG^SF^ as an adjuvant elicited an exceptional high antibody titer and a robust T cell response, which were better than the group using aluminium hydroxide as a adjuvant. These results highlighted the adjuvant potential of STING agonist in SARS-CoV-2 vaccine preparation for the first time.

## Main Text

Being a potent immunostimulator, cyclic dinucleotides (CDNs) could activate the STING (STimulator of INterferon Genes) pathway, leading to the production of type I interferon (IFN β) and other pro-inflammatory cytokines.^1,2^ Type I interferon plays a vital role in regulating the cross-priming of CD8+T cells.^3^ Locating on the endoplasmic reticulum of antigen-presenting cells (APCs) and B cell, STING protein could bind cyclic 2’,3’-GAMP produced by cyclic GMP-AMP synthase (cGAS) or cyclic di-GMP (CDG), cyclic di-AMP (CDA) and 3’3’-GAMP from bacteria.^4,5^ STING pathway serves as a central mediator of antitumor/antivirus immunity and has been an attractive target for cancer immunotherapy and auto-immune diseases treatment. Intratumoral administration of phosphorothioated CDA alone or with anti-PD-1 (Keytruda), anti-CTLA-4 (Ipilimumab) are being studied in phase Ib clinical trials by Novartis and Merck (NCT02675439, NCT03010176).^4^

The coronavirus disease in 2019 (COVID-19) caused by the infection of SARS-CoV-2 is still in a pandemic around the world, which has resulted in more than 12,000,000 cases and more than 520,000 deaths. To control the ongoing global pandemic, the research and development of therapeutic and prophylactic interventions like Remdesivir (Veklury) and vaccine are being speeded up.^6^ As an effective strategy to contain the large-scale transmission, SARS-CoV-2 vaccines have been placed in great hopes. SARS-CoV-2 is found to be a β-coronavirus and depends on the interaction between its surface envelope-embedded spike (S) glycoproteins and human angiotensin-converting enzyme 2 (hACE2) receptors to initiate virus entry, just like SARS-CoV.^7^ Therefore, to block the infection, S protein has been recognized as a key antigen for vaccine preparation.^8^

Inactivated virus vaccine, recombinant S protein vaccine, DNA/RNA vaccine, and recombinant adenovirus (S protein gene) vectored vaccine are the major COVID-19 vaccine types studied in preclinical and clinical trial.^9^ Except vectored vaccine, other types, especially inactivated and recombinant protein vaccine, need the help of adjuvant like aluminium hydroxide to elicit a robust and durable immune responses while reducing antigen doses and injection times. Though approving as an adjuvant by FDA, aluminium hydroxide usually induces a Th2-dominant response and hardly elicit cellular immunity.^10^ While, Dong and Sette et al. had respectively identified a high percent (70% CD8+T, 100% CD4+T) SARS-CoV-2-specific cytotoxic T cells in COVID-19 convalescent patients, correlating with antibody titers.^11,12^ This indicates a key role of adaptive immunity in clearing virus-infected cells and reducing inflammation.^13^ Considering great efficacy of CDNs in producing type I IFN for T cell response, we decide to explore the adjuvant effect of CDNs on SARS-CoV-2 recombinant S protein vaccine for the first time and hope to provide a novel choice for coping with emerging pandemic (Fig. 1a).

**Fig. 1.**
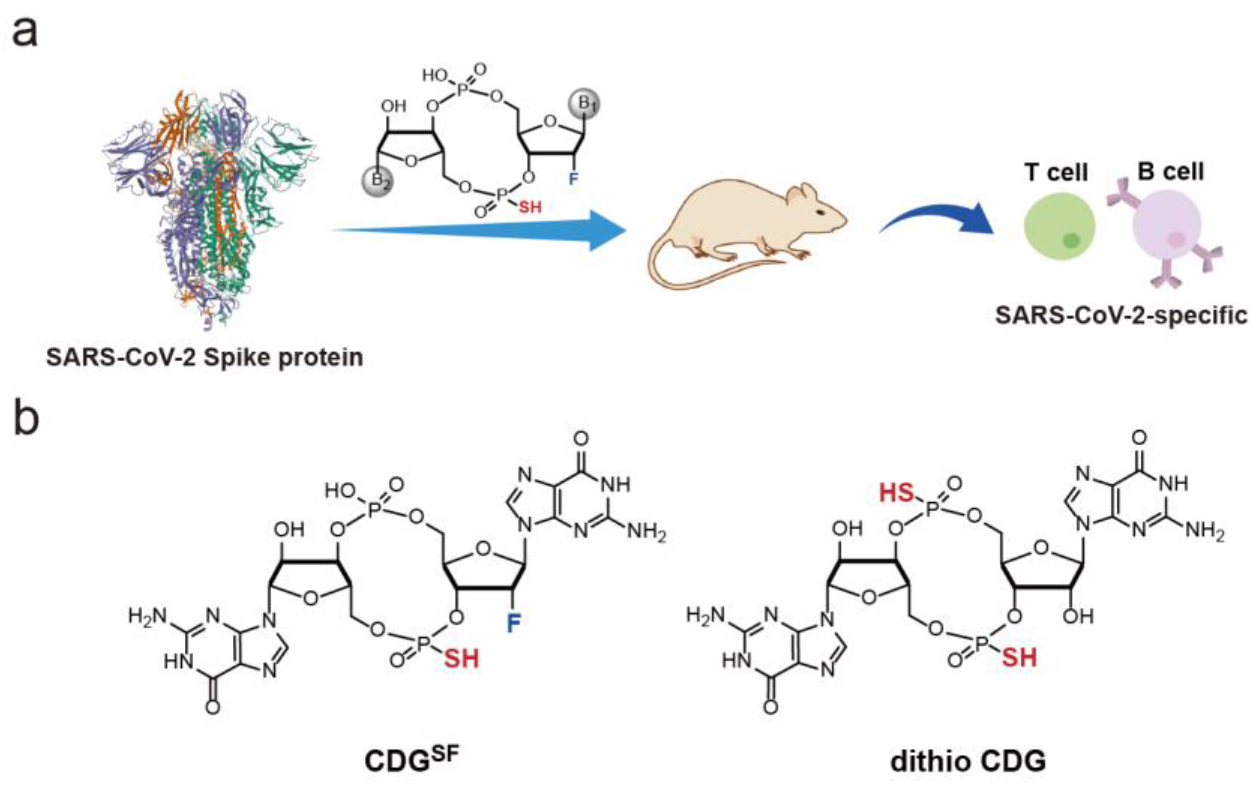
(a) The SARS-CoV-2 vaccine adjuvant based on a modified STING agonist. (b) Chemical structure of CDG^SF^ and dithio CDG. B1/B2: base.

Despite the potency of CDNs in immunotherapy and adjuvant, the negative charges, instability and hydrophilicity have greatly impeded their biomedical applications.^4^ Two phosphate groups in CDNs restrict their entry into the cell cytosol to interact with STING protein. And the degradation of CDNs by phosphoesterase also reduces their stimulating efficiency. To overcome the drawbacks, we decided to modify the CDNs structure with fluorine and phosphorothioate.

Here, we chose CDG as a model CDNs. To improve CDG stability, we have synthesized a novel CDG analogue CDG^SF^ for the first time which replaced one 2’-OH with F atom and one phosphate diesters with phosphorothioate diesters at the same side (Fig. 1b). CDG^SF^ showed enhanced activity compared with phosphorothioated CDG (dithio CDG, Fig. 1b). Besides, recombinant S protein immunization with CDG^SF^ as an adjuvant elicited a robust humoral (IgG) and cellular (T cell) response, which was better than aluminium hydroxide group.

### Synthesis of CDG analogue CDG^SF^

The CDG analogue CDG^SF^ was synthesized based on Jones and co-workers’ one flask solution-phase strategy.^14–16^ After a few attempts, some conditions and raw materials had been adjusted as shown in synthetic route (Scheme 1). The commercial available 2’-F guanosine phosphoramidite, **D3**, was adopted as the second portion to couple with deprotected guanosine **D2**. To obtain the mono-phosphorothioate modification, sulfurization with 3-((dimethylaminomethylidene)-amino)-3H-1,2,4-dithiazole-5-thione (DDTT) was conducted after coupling. Following cyclization with 2-chloro-5,5-dimethyl-1,3,2-dioxaphosphorinane-2-oxide (DMOCP), the compound was oxidized with iodine to form the second phosphodiester (**D6**). Finally, the target molecule CDG^SF^ was acquired through de-protection and crystallization, with an overall yield of 40%.

**Scheme 1.**
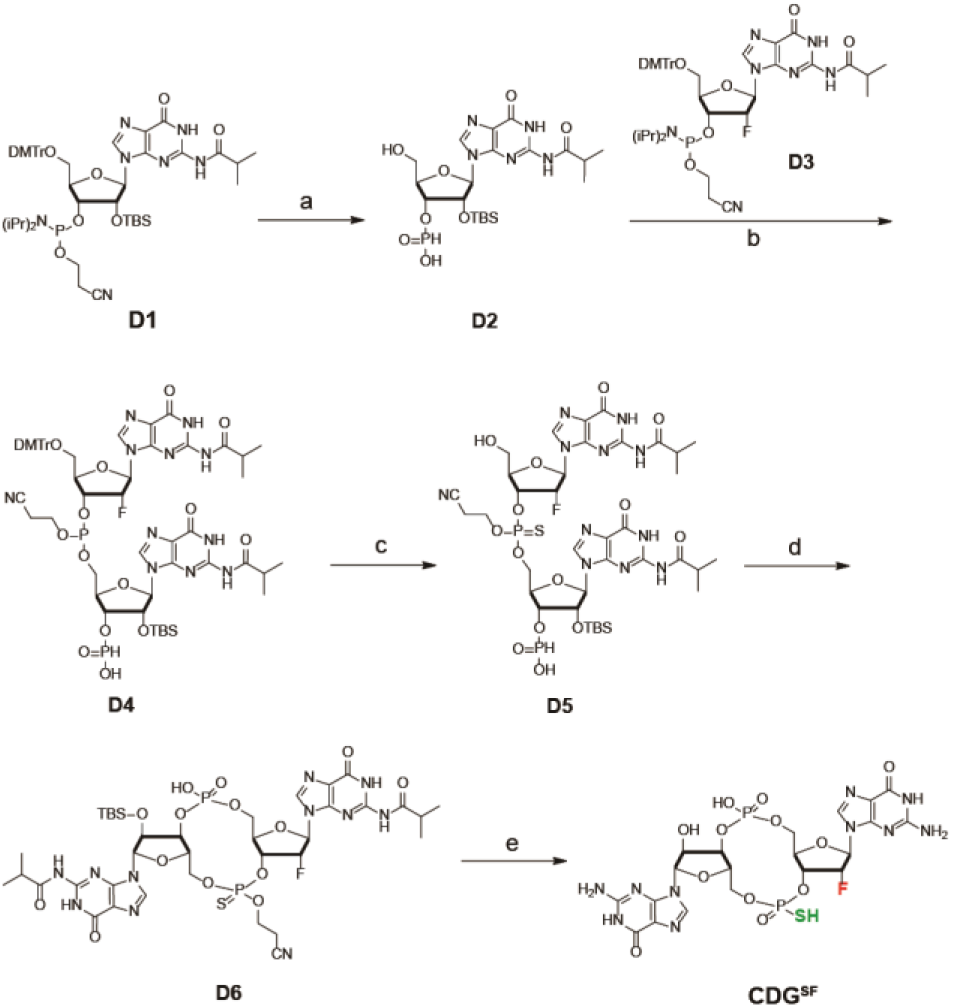
Chemical synthesis route of CDG^SF^. a i) pyridine-TFA, H_2_O; ii) t-BuNH_2_; iii) DCA (dichloroacetic acid), H_2_O; b pyridine; c i) DDTT; ii) DCA, H_2_O; d i) DMOCP; ii) I_2_, H_2_O; e i) t-BuNH_2_, ii) CH_3_NH_2_, iii) Et_3_N-HF.

### CDG^SF^ could efficiently activate macrophages in vitro

After finishing the synthesis, we were curious about the activity of CDG^SF^ which had not been studied before. We adopted J774A.1 cells (a STING-expressing mouse macrophage) as a model cell line, since the macrophage is a critical immune cell for the antitumor and adjuvant effect of CDNs. Considering that phosphorothioated CDNs is the preferential form for clinical researches, ^17^ we adopted dithio CDG (Fig. 1b) as a control group. Because of poor penetrability mentioned above, we added the agonists to cells with or without the pre-treatment of a transfection reagent (see methods for details). After incubation, the cells were labelled with anti-CD86 antibody for flow cytometry analysis. As illustrated in Fig. 2, both of CDG^SF^ and dithio CDG with transfection remarkably upregulated the expression level of activation marker co-stimulatory molecule CD86. More importantly, CDG^SF^ induced higher CD86 level compared to dithio CDG either with or without the transfection. The better performance of CDG^SF^ could be owed to the fluorine modification which brought CDG^SF^ with improved liposolubility and stability.^18^ These results indicated that CDG^SF^ was capable of stimulating STING pathway efficiently and might be a better clinical choice relative to dithio CDG. Besides, the poor results of dithio CDG and CDG^SF^ without transfection further highlighted the importance and urgency of developing the delivery method (Fig. 2).

**Fig. 2.**
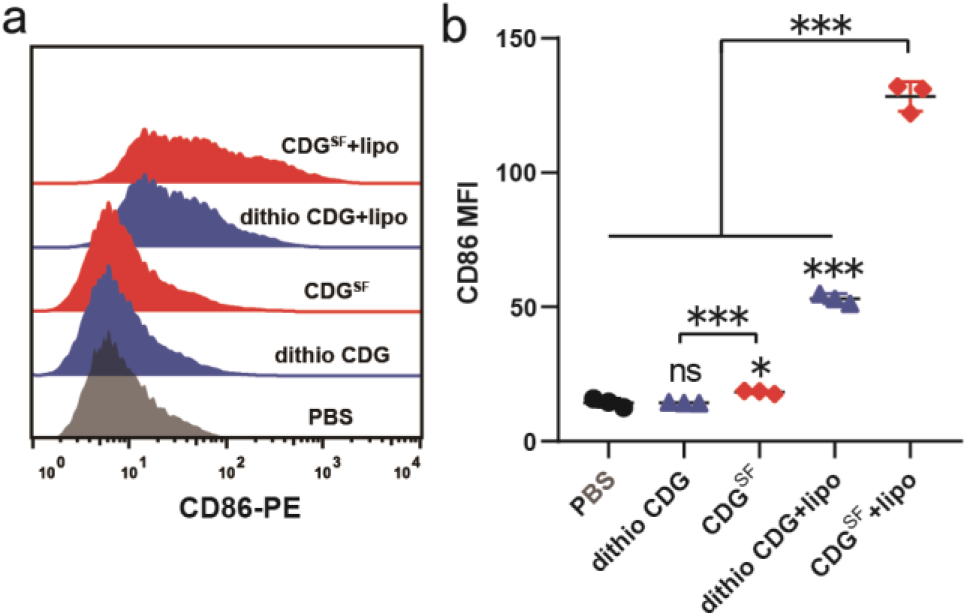
Expression of CD86 on J774A.1 cells stimulated by CDG^SF^ and dithio CDG with or without liposome transfection (10 μM). PBS was used as the negative control. (a) Flow cytometry and (b) CD86-PE fluorescence intensity. Stimulation indexes were displayed as the mean fluorescence intensity (MFI). Results are shown as mean ± SD of three separate experiments, ns: no significant difference, *p<0.05, ***p<0.001 by Student’s t-test.

### Immunization of SARS-CoV-2 Spike protein with CDG^SF^ induced a robust T cell response

To explore the adjuvant effect of CDG^SF^ on SARS-CoV-2 vaccine, we chose recombinant Spike S1+S2 extracellular domain (ECD) glycoprotein (sequence: YP_009724390.1) as a model antigen. Besides, Spike (S) protein plus aluminium hydroxide (Alhydrogel® adjuvant 2%, 100 μg per mouse) was adopted as a control group. Immunizations on Babl/c mice were performed for three times biweekly (Fig. 3a). One week after last injection, sera and spleen samples were harvested. SARS-CoV-2 S protein-specific T cell response was assessed through IFN-γ enzyme-linked immunospot assay (IFN-γ ELISPOT, see methods for details). Splenocytes were stimulated with 50 μg/mL S protein for 36 h before forming IFN-γ spot. As shown in Fig. 3b, the number of IFN-γ-secreting S protein-specific T cells in CDG^SF^ group was much higher than Alum and S protein group. As expected, Alum adjuvant contributed the limited promotion to the cellular response of S protein. Besides, the spot numbers were corroborated by mouse spleen weight of each group (Fig. 3c). This result proved that CDG^SF^ could be an excellent adjuvant to significantly improve the SARS-CoV-2-specific T cell responses, prior to Alum.

**Fig. 3.**
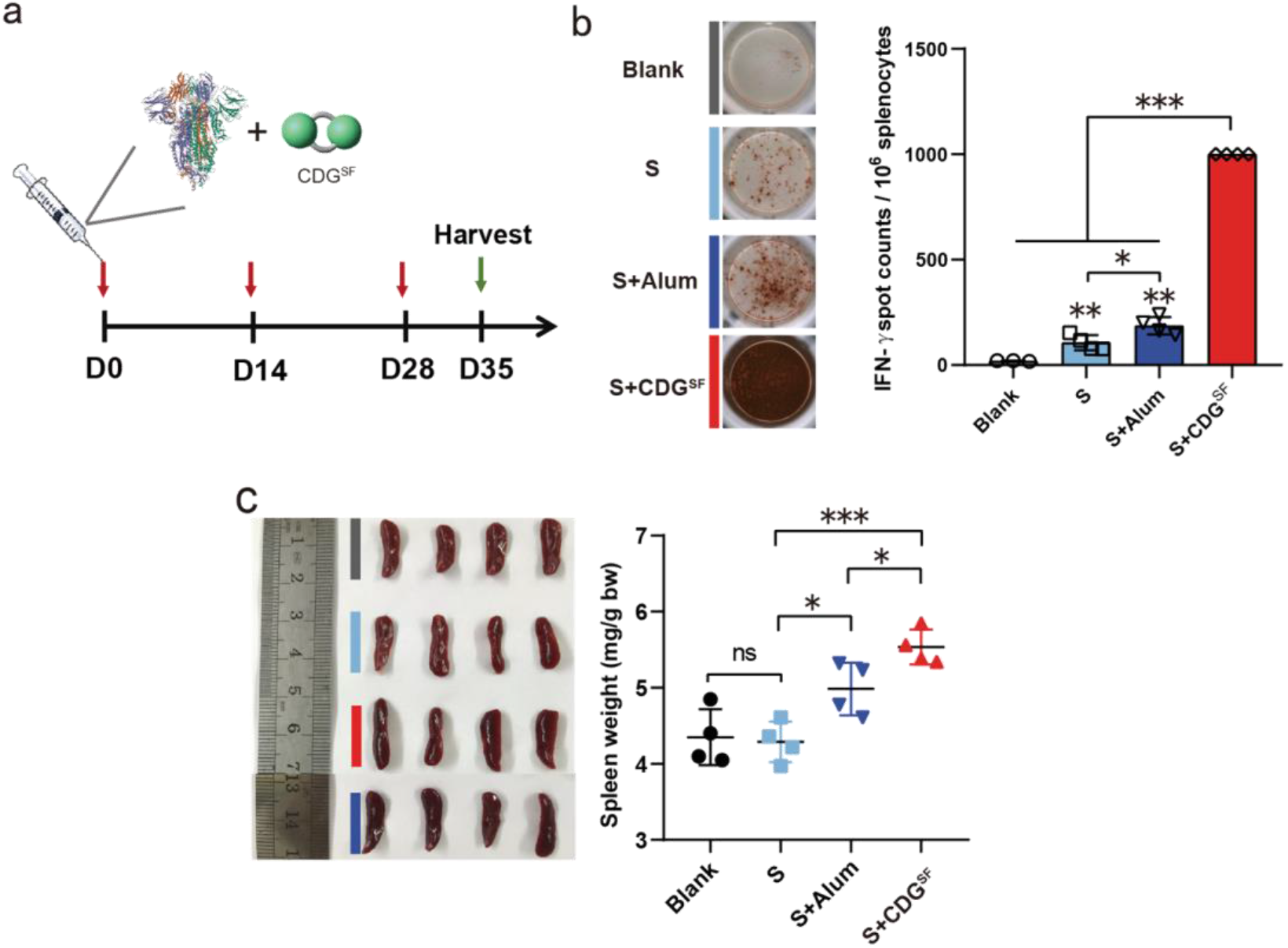
Immune analysis of CDG^SF^ as an adjuvant for SARS-CoV-2 vaccine. (a) Immunization scheme of the vaccine group: blank, S, S+Alum and S+CDG^SF^. (b) IFN-γ ELISPOT test of splenocytes and (c) spleen weight data from immunized mice. ns: no significant difference, *p<0.05, **p<0.01, ***p<0.001 by Student’s t-test.

### Immunization of SARS-CoV-2 Spike protein with CDG^SF^ elicited an exceptionally high IgG titers

As to humoral response, we adopted enzyme linked immuno-sorbent assay (ELISA) to analyse the S protein-specific IgG titers with sera from immunized mice. The plate was precoated with 0.1 μg/well S protein. As illustrated in Fig. 4a, S protein plus CDG^SF^ immunization elicited an exceptionally high SARS-CoV-2-specific IgG titer (endpoint titer up to 819,200), close to Qin’s Inactivated SARS-CoV-2 vaccine and higher than IgG titers (about 20,000) in recovered COVID-19 patients.^19^ While, Alum group did not exhibit the adjuvant effect on S protein compared with S protein alone, suggesting the need of higher aluminium hydroxide dose. The titers data demonstrated that CDG^SF^ could also notably enhance S protein-specific humoral response.

**Fig. 3.**
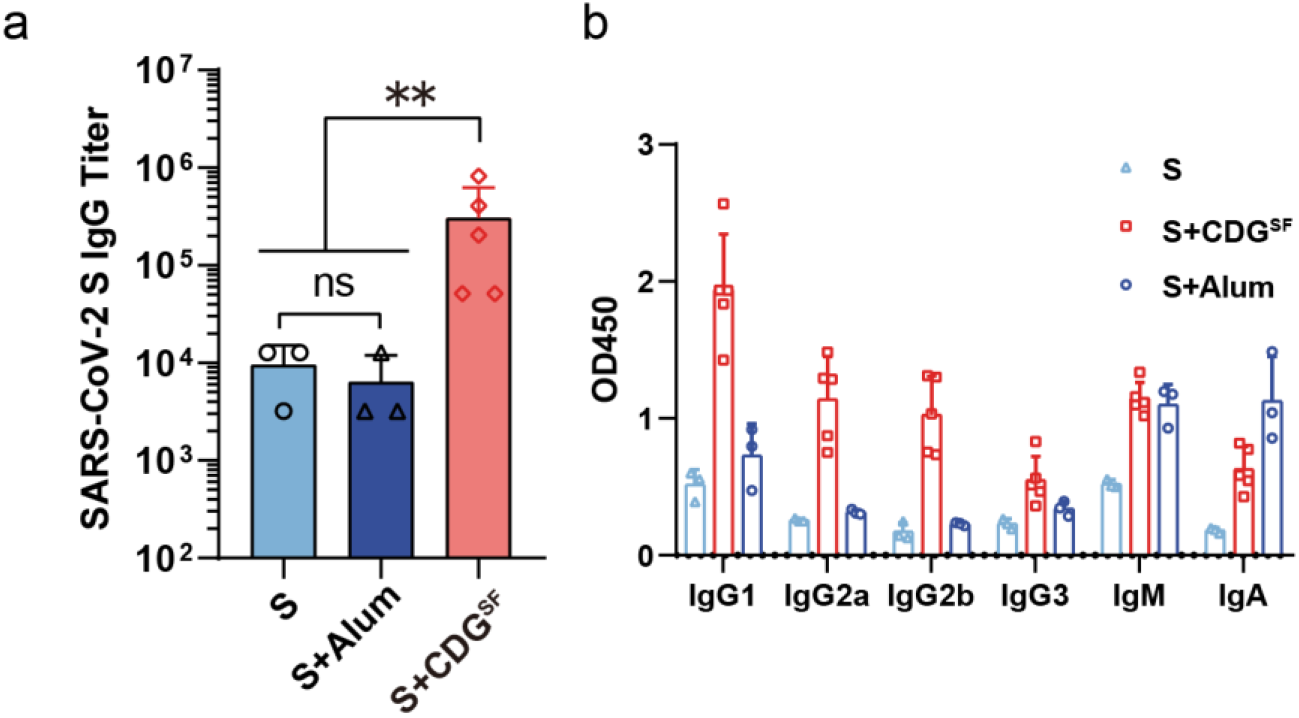
SARS-CoV-2 specific antibody titers (a) and isotypes (b) evaluation in the antisera of each vaccine group. ns: no significant difference, **p<0.01 by Student’s t-test.

To further compare the relative intensity of cellular and humoral response induced by CDG^SF^ immunization, we detected antibody isotypes distribution in sera (Fig. 4b, dilution 1:12800). In general, the level of IgG1 isotype is related to humoral response (Th2) and the generation of IgG2a reflects the T cell activation (IFN-γ, Th1). In anti-sera of CDG^SF^ group, the amount of IgG1 was relatively more than IgG2a, IgG2b and IgM, along with minimal levels of IgG3 and IgA. The level ratio (2:1) of IgG1 and Ig2a revealed that the combination of S protein and CDG^SF^ generated a balanced cellular and humoral (Th1/Th2) response. In Alum group, the higher level of IgM over IgG1 and IgG2a indicated an inefficient immune activation. The isotype results again highlighted the adjuvant potential of CDG^SF^ in SARS-CoV-2 vaccine preparation.

## Conclusion

In brief, we have designed and synthesized a novel CDN CDG^SF^ which exhibited better STING activity over dithio CDG. Besides, we proved for the first time that CDG^SF^ could act as an excellent adjuvant to notably improve the S protein-specific T cell and IgG titer level of SARS-CoV-2 vaccine, overcoming the drawbacks of aluminium hydroxide. These results also suggest that STING agonists might provide a great and general adjuvant choice for multiple kinds of SARS-CoV-2 vaccines including inactivated virus vaccine, recombinant RBD protein vaccine, peptide vaccine and DNA/RNA vaccine.

This work was supported by National Key R&D Program of China (2019YFA0904200, 2018YFA0507600) and National Natural Science Foundation of China (21672126). The synthesis of CDNs compounds and adjuvant application of STING agonist in SARS-CoV-2 vaccine are applying for China patents (Patent No 202010663965.4, 202010411675.0 and 202010694692.X).

## Methods

### Synthesis of CDG^SF^

The CDG^SF^ was synthesized based on the one-flask synthesis strategy by Jones et al with some adjustments.^14,15 1^H NMR (400M, D_2_O) δ 8.26–7.71 (m, 2H), 6.38–5.87 (m, 2H), 5.86–5.45 (m, 2H), 4.58–4.32 (m, 4H), 4.07 (d, *J* = 11.1 Hz, 2H). ^31^P NMR (400M, D_2_O) δ 55.35, 54.96, −1.05. ^19^F NMR (400M, D_2_O) δ −122.42, −130.51. ESI-HRMS (negative mode): C_20_H_22_FN_10_O_12_P_2_S^−^ [M-H]^−^ calculated 707.0604; found 707.0606.

### Evaluation of macrophage activation in vitro using J774A.1 cell line

J774A.1 cells (mouse monocyte macrophage cell line) were cultured in DMEM (dulbecco’s modified eagle medium) containing 10% fetal bovine serum (FBS) at 37°C, 5% CO_2_. After harvest, the cells were planted on 24-well culture plates with a density of 5×10^5^ cells/well and cultured overnight. Then, 10 μM of the compounds were added respectively and incubated for 18 h. For the transfection group, Lipofectamine® 3000 was used as a transfection reagent and conducted according to manufacturer’s protocol. Samples were mixed with DMEM as a A solution (150μL). Lipofectamine® 3000 (4.5 μL) was mixed with DMEM as a B solution (150 μL). 5 min later, solution A was added to the solution B. Waiting for 15 min, the mixture was added dropwise to 24-well plate at a final concentration of 10 μM and incubated for 18h. Then, cells were harvested and strained with mouse anti-CD86-phycoerythrin antibodies (BD Pharmingen, dilution 1/200) at ice for 1h. After washing, the cells were analyzed on BD Calibur flow cytometry.

### SARS-CoV-2 vaccines immunization

6-8 week old Babl/c mice (4-5 mice per group, female) were separately subcutaneously vaccinated with SARS-CoV-2 S protein 5 μg/mouse, CDG^SF^ 20 μg/mouse, Alhydrogel® adjuvant 2% 100 μg/mouse. SARS-CoV-2 Spike protein (S1+S2 ECD, gene: YP_009724390.1) was purchased from Sino Biological Inc. Alhydrogel® adjuvant 2% was purchased from InvivoGen. Immunizations were conducted for three times biweekly. Antisera and spleens were collected one week after the last administration. Mice used in the experiments were raised in Animal Facility of Center of Biomedical Analysis in Tsinghua University and treated in compliance with the animal ethics guidelines. The animal protocol (approval number: 16-LYM2) was approved by Institutional Animal Care and Use Committee (IACUC) of Tsinghua University. Animal Facility of Center of Biomedical Analysis in Tsinghua University has been authenticated by Association for Assessment and Accreditation of Laboratory Animal Care (AAALAC).

### IFN-γ enzyme-linked immunospot assay (IFN-γ ELISPOT)

IFN-γ ELISPOT Kit was purchased from Dakewe Biotech Co., Ltd. Spleens from immunized mice were grinded and filtered through a 40 μm cell strainer. And red blood cells (RBCs) were lysed using lysis buffer for RBC. Splenocytes were counted and added to 96-well kit at 1000,000 amount/ 100 μL per well. Then SARS-CoV-2 S protein was added to the well (final concentration: 50 μg/mL) and cells were stimulated for 36 h. The spot forming procedure was performed according to the Kit instruction.

### SARS-CoV-2-specific antibody titers

1 μg/mL SARS-CoV-2 S protein in coating buffer (0.1M NaHCO_3_ solution, pH=9.6) was added to high-binding 96-well ELISA plate (Costar 3590, 100 μL/well). After incubation for 12h at 4°C and washing with PBST solution (0.05% Tween in PBS buffer), the wells were blocked by 0.25% gelatin PBS solution for 3h at room temperature. After washing with PBS and PBST, the diluted antisera (1:200) was added to each well (100 μL per well) and incubated for 1.5h at 37°C. After washing again, diluted rabbit anti-mouse IgG-Peroxidase antibodies (1/2000 dilution, Sigma) were added to each well (100 μL per well) and incubated for 1h at 37°C. After washing and spin-drying, 3,3’,5,5’-Tetramethylbenzidine (TMB) was added to plate (200 μL per well) and incubated for 4 min and then stopped by 2 M H_2_SO_4_ (50 μL per well). Optical absorption was measured at wavelength of 450nm and antibody titer was defined as highest dilution yielding an optical absorption of 0.1 or greater over that of negative control antisera.

### SARS-CoV-2-specific antibody isotypes

96-well ELISA plate was coated with SARS-CoV-2 S protein according to the procedure described above. The antisera were diluted to 1:12800 and added to each well, incubating for 1.5h at 37°C. After washing with PBS and PBST, isotype antibodies IgG1, IgG2a, IgG2b, IgG3, IgA and IgM (anti-mouse antibodies from goat, Sigma) were diluted to 1:1000 and added to each well (100μL per well). After incubation for 1.5 h at 37°C and washing, 200μL TMB substrate described above was added to plate, incubated for 4 min and stopped by 2 M H_2_SO_4_ (50 μL per well). Optical absorption was also measured at wavelength of 450 nm.

